# Fragile X Premutation rCGG Repeats Impairs Synaptic Growth and Synaptic Transmission at *Drosophila* larval Neuromuscular Junction

**DOI:** 10.1101/2020.10.22.349928

**Authors:** Sajad Ahmad Bhat, Adil Yousuf, Zeeshan Mushtaq, Vimlesh Kumar, Abrar Qurashi

**Author notes:** **Corresponding Author:** Abrar Qurashi^1^.

## Abstract

Fragile X-associated tremor/ataxia syndrome (FXTAS) is a progressive neurodegenerative disease manifesting in the premutation (PM) carriers of the *FMR1* gene with alleles bearing 55-200 CGG repeats. The discovery of a broad spectrum of clinical and cell developmental abnormalities among PM carriers with or without FXTAS, and in model systems suggests that neurodegeneration seen in FXTAS could be the inevitable end-result of pathophysiological processes set during early development. Hence, it is imperative to trace early pathological abnormalities. Our previous studies have shown that transgenic *Drosophila* carrying human-derived fragile X premutation-length CGG repeats are sufficient to cause neurodegeneration. Here, we used the same transgenic *Drosophila* model to understand the effects of fragile X premutation-length CGG repeats on the structure and function of the developing nervous system. We show that presynaptic expression of the premutation length CGG repeats restricts synaptic growth, reduces the number of synaptic boutons, leads to aberrant presynaptic varicosities, and impairs synaptic transmission at the larval neuromuscular junctions (NMJs). The postsynaptic analysis shows both glutamate receptor and subsynaptic reticulum proteins are normal. However, a high percentage of boutons show the reduced density of Bruchpilot protein, a key component of presynaptic active zones required for vesicle release. The electrophysiological analysis shows a significant reduction in the quantal content, a measure of total synaptic vesicles released per excitation potential. Together these findings endorse that synapse perturbation caused by rCGG repeats mediate presynaptically during larval NMJ development.

## Introduction

The Fragile X Mental Retardation 1 (*FMR1*) gene normally harbors a highly polymorphic trinucleotide repeat sequence (CGG) within its 5’ untranslated region (5’UTR). The normal allele of the *FMR1* gene typically has 5 to 40 CGG repeats. Abnormal alleles include the full-mutation (> 200 CGG repeats), premutation (55-200 CGG repeats), and the gray zone mutation (45-54 CGG repeats). Carriers of full mutation lead to Fragile X Syndrome (FXS), the most common inherited form of neurodevelopment and Intellectual Disability (ID) disorder, and occur in 1 in 4000 to 1 in 7000 people (Kremer et al. 1991, Verkerk et al. 1991, Hagerman and Hagerman 2002, Colak et al. 2014). On the other hand, PM carriers account for a variety of phenotypes that are found frequently in the population with an estimated prevalence of 1:259 in females and 1:813 in males (Dombrowski et al. 2002, Rousseau et al. 1995). A proportion of these PM carriers, about 40% of males and 16% of females develop a progressive neurodegenerative disorder termed fragile X-associated tremor/ataxia syndrome (FXTAS) usually after the fifth decade of life (Hagerman and Hagerman 2015, Jacquemont et al. 2004). Clinically, FXTAS presents with intention tremor, gait ataxia, and other features including parkinsonism, cognitive defects, brain atrophy, and white matter abnormalities on MRI (Jacquemont et al. 2003, Hagerman and Hagerman 2015). Neuropathologically, FXTAS is distinguished by the presence of large eosinophilic nuclear inclusions in neurons, astrocytes, and in the spinal column as well as peripheral tissues (Greco et al. 2006, Ariza et al. 2016, Greco et al. 2002). These inclusions contain the expanded *FMR1* messenger RNA (mRNA), ubiquitin, as well as several other proteins that aggravate cellular dysregulation (Iwahashi et al. 2006).

One unique molecular signature of the fragile X PM allele is that the level of *FMR1* mRNA is significantly elevated, however, *FMR1* encoded protein (FMRP) paradoxically remains relatively unchanged. So the neurodegenerative phenotypes associated with FXTAS are suspected of being caused by a gain of function in fragile X PM rCGG-repeat RNAs (Tassone et al. 2000c, Tassone et al. 2000b, Tassone et al. 2000a, Tassone, Hagerman and Hagerman 2014). Model systems developed to mimic the molecular, cellular alterations and clinical symptoms of FXTAS have proposed several mechanisms of pathogenicity that include RNA-mediated toxicity and repeat-associated non-AUG (RAN) mediated proteotoxicity (Jin et al. 2003, Jin et al. 2007, Kong et al. 2017, Qurashi et al. 2011, Sofola et al. 2007, Glineburg et al. 2018). The RNA gain-of-function, however, seems to play a major role in rCGG mediated pathogenesis (Jin et al., 2003, Sofola et al., 2007a, Jin et al., 2007, Sellier et al., 2010). It is proposed that overproduced rCGG repeats in FXTAS sequester specific RNA-binding proteins and reduce their ability to do their normal cellular functions, thereby contributing significantly to the pathology of this disorder (Galloway et al. 2014, Jin et al. 2007, Qurashi et al. 2011, Sofola et al. 2007, Tan et al. 2012a, Tan et al. 2012b, Sellier et al. 2010, Sellier et al. 2013).

The disease initially identified as neurodegenerative was complicated with a broad spectrum of clinical abnormalities among PM carriers, which are apparent both at an early age and most likely in a larger cross-section of carriers than those who develop FXTAS later in life (Ariza et al. 2016). The PM carrying children were found to have elevated levels of *FMR1* mRNA and comorbid for neurodevelopmental disorders like autism, attention deficit hyperactivity disorder (ADHD) (Hagerman et al. 1996, Tassone et al. 2000, Goodlin-Jones et al. 2004). A study of 43 male subjects including 27 PM carriers showed a significant co-occurrence of Autism Spectrum Disorder (ASD) and Attention Deficit Hyperactivity disorder (ADHD) in both proband and non-proband PM carriers (Farzin et al. 2006). Chonchaiya and colleagues also observed an increased occurrence of autism and ASD features in PM carriers with respect to the sibling control (Chonchaiya et al. 2012). A national survey of families with FXS children, which included 256 PM subjects revealed increased cases of developmental abnormalities in PM carriers (Bailey et al. 2008).

Experimental investigations focused on early abnormalities have shown mitochondrial dysfunction in fibroblasts and brain samples of PM carriers with and without FXTAS (Alvarez-Mora et al. 2017, Ross-Inta et al. 2010). Embryonic fibroblasts from the KI mouse have shown abnormal lamin A/C architecture with loss of ring-like nuclear staining (Garcia-Arocena et al. 2010). Abnormalities during embryonic development included neuronal migration defects, reduced dendrite length, and dendritic complexity (Chen et al. 2010). Corroborating studies from the second KI mouse have shown dendritic morphology abnormalities in cultured hippocampal neurons (Qin et al. 2011, Cao et al. 2012, Hunsaker et al. 2009). The PM KI mice also manifested behavioral and cognitive deficits. In behavioral studies, there were progressive deficits in spatial processing (but no motor involvement) in mice as young as 12 weeks (Hunsaker et al. 2009, Berman and Willemsen 2009). Also, female PM carriers aged between 21-42 years showed early neurocognitive impairments than the motor/cognitive impairments associated with FXTAS (Goodrich-Hunsaker et al. 2011).

The *Drosophila* disease model of FXTAS has provided experimental evidence for RNA mediated pathogenic mechanisms. This model has generally been employed for genetic modifier screens because of its convenient phenotypic readouts in the eyes. Although several important proteins/pathways have been identified through this system (Jin et al. 2003, Jin et al. 2007, Qurashi et al. 2011, Sellier et al. 2017, Todd et al. 2013), the eye phenotype, however, represents a late stage when the neurons have much deteriorated. The larval neuromuscular junctions (NMJs) are large synapses that are easily accessible to investigate. They offer leeway for a detailed temporal and functional analysis making the extrapolations more accurate (Prokop 2006, Ehmann, Owald and Kittel 2018). Hence, we sought to explore the early pathological consequences of fragile X PM rCGG repeats on the NMJ structure and function using the *Drosophila* disease model of FXTAS.

We show that fragile X PM rCGG repeats impairs larval locomotion and causes a significant change in the overall morphology of synapses at larval Neuromuscular Junction. Total synapse length including the number of branches and boutons were significantly reduced than controls. However, boutons, particularly at the terminals were larger than normal. The aberrant bouton morphology precedes with a significant decrease in the density of Bruchpilot protein at active zones, the sites of vesicular release. In accord, rCGG repeats compromise synaptic transmission. Together we show that expression of fragile X PM rCGG repeats exacerbate synaptic growth and transmission when repeats are specifically expressed in the presynaptic compartment of synapses during larval NMJ development.

## Results

### Neuronal Expression of Fragile X Premutation riboCGG (rCGG) Repeats Induce Defects in Larval Locomotion

FXTAS is long considered to be late onset neurodegenerative disease (Hagerman 2013). However, recent studies suggest FXTAS could be the end-stage of a life-long process of neuronal deregulation (Hagerman 2012). On this premise, we sought to examine early neuronal phenotypes in transgenic flies with both moderate (pUAST-(CGG)_60_-EGFP), or strong (pUAST-(CGG)_90_-EGFP) transgene expressions (Jin et al. 2003, Jin et al. 2007, Qurashi et al. 2011). We evaluated transgene expression levels through RT-PCR (data not shown), and by directing transgene expression to several tissues. The transgenic line with the pUAST-EGFP vector alone served as control.

We first analyzed the effects of the fragile X PM rCGG repeats in embryos. We directed transgene expression to all developing cells of the peripheral and central nervous system using the *elav-GAL4* driver. At 25 °C, strong expression of r(CGG)_90_, causes lethality primarily during embryonic development before larval formation (Jin et al. 2003). However, moderate r(CGG)60 expression produced viable and full-term embryos. We labeled these embryos with anti-FasII, and anti-BP102 antibodies, which recognize different subsets and subcellular compartments of neurons in the central and in the peripheral nervous system, respectively. A high percentage of these embryos did not display any obvious or gross axon morphology defects (97% of the CGG embryos (n=150)). However, during the larval stage, they exhibited a severely compromised locomotion. In accord with *ok6-GAL4,* motor neurons specific driver, expression of r(CGG)_90_ repeats displayed a characteristic larval tail-flip phenotype where the posterior body segments flip upwards after each peristaltic muscle contraction (Figure 1A). To determine the specificity of this phenotype, we performed a systematic assay based on locomotion activity. In this assay, wandering third instar larvae were allowed to acclimatize on the agarose plates and were monitored over time by measuring the number of grids (1 sq.mm) crossed by individual third-instar within a 60 sec time window over a test period of 180 seconds. As shown in Figure 1B the locomotion of both r(CGG)60 and r(CGG)90 was significantly reduced compared to controls, (Control vs (CGG)_90_; 0.55 ± 0.02 versus 0.13 ± 0.01 mm/sec; mean ± SEM; p < 0.0001). Importantly, compared to controls we did not see a significant difference in locomotion when similar repeats were driven with the *mhc-GAL4* driver, a postsynaptic driver. This indicates that locomotion impairments could originate more in neurons than in muscle cells.

**Figure 1:**
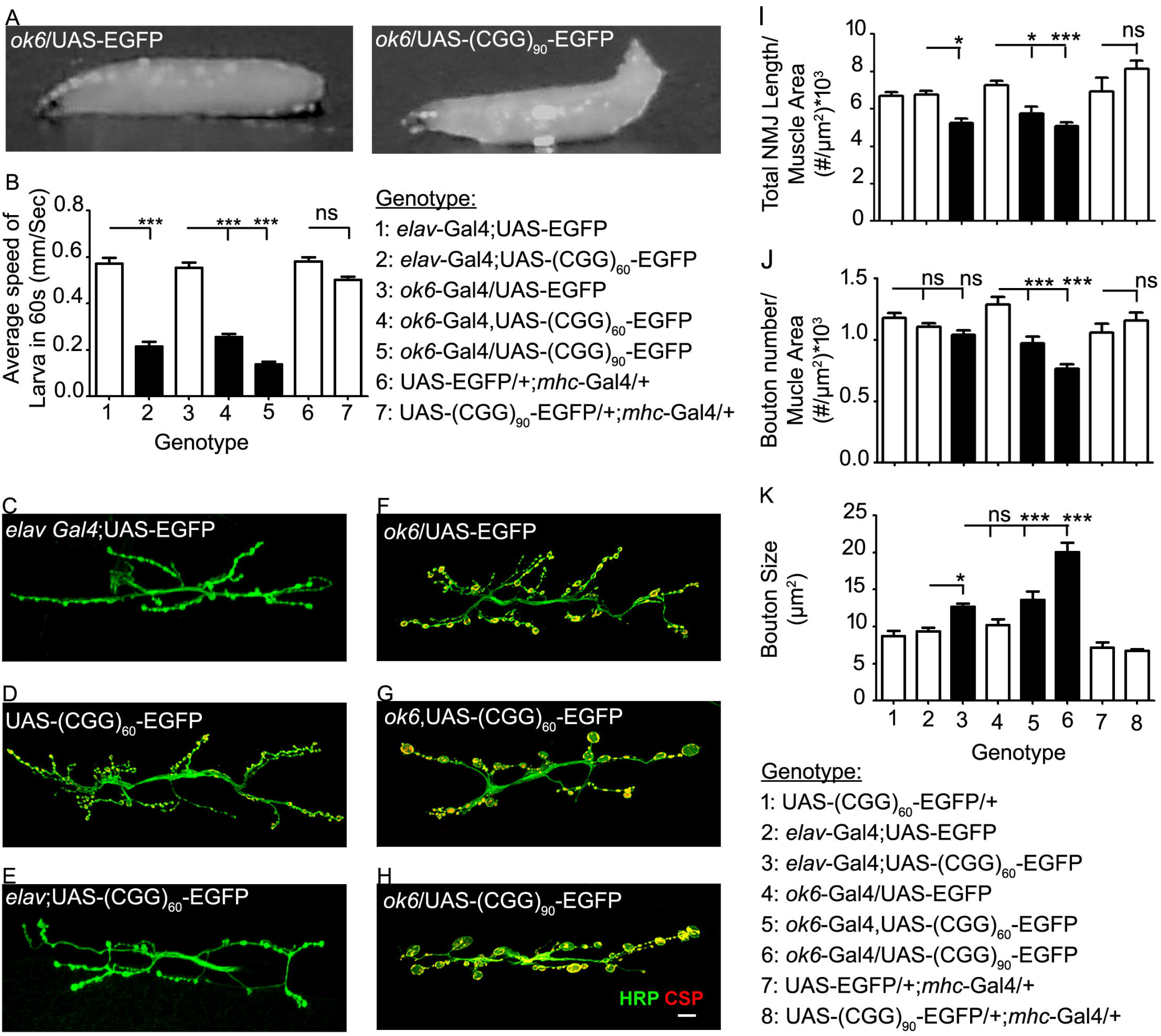
Expression of premutation rCGG repeats in neurons causes locomotion activity and larval NMJ defects. (A) Representative larvae showing the characteristic tail-flip phenotype in rCGG expressing larvae (*ok6-GAL4/*+; UAS-(CGG)90-EGFP/+) at 25 °C while the control (*ok6-GAL4*/+) does not show this behavior. (B) Histograms comparing crawling speed where the numerical values in the bars show the number of larvae quantified. Student’s t-test was used for statistical analysis. *** represents p<0.0001. Error bars represent mean ± s.e.m. (C) Representative confocal images of wandering third instar larval neuromuscular junction muscle 6/7 of *elav-GAL4;* UAS-EGFP; (D) UAS-(CGG)60-EGFP/+; (E) *elav-GAL4;* UAS-(CGG)60-EGFP at 25°C (F) *ok6*-*GAL4*/UAS-EGFP; (G) *ok6-GAL4,* UAS-(CGG)60-EGFP; (H) *ok6-GAL4* /UAS-(CGG)90-EGFP at 25°C co-labeled with antibodies against cysteine string protein (CSP) in red and horseradish peroxidase (Hrp) in green. The Scale bar represents 10μm. Histograms show (I) total NMJ length normalized to the surface area of their corresponding muscle (length/area in μm) × 10, (J) number of boutons per unit muscle area (number/area in μm^2^) × 10^3^, and (K) bouton size in μm^2^. *** represents p<0.0001 and ** represents p<0.001. Error bars represent mean ± s.e.m. Student’s t-test was used for one to one analysis and One-way ANOVA with Tukey post-hoc was used for multiple comparisons.

### Neuronal Expression of Fragile X Premutation riboCGG (rCGG) Repeats Induce Defects in the Morphology of Larval Neuromuscular Junctions (NMJs)

The fact that neuronal expression of rCGG repeats leads to compromised locomotion in larvae prompted us to examine their neuromuscular morphology. We performed open-book preparations of larval body walls (Prokop 2006, Menon, Carrillo and Zinn 2013), and used anti-horseradish peroxidase (HRP) to label the neuronal membrane of motor axons. In larvae expressing moderate (CGG)_60_-EGFP transgene with the *elav-GAL4* driver, motor axons emanating from ventral nerve cord (VNC) followed their normal stereotyped pattern of innervations, however, synaptic arbors appeared altered in their gross morphology. Based on this finding, we systematically examined NMJs for type-Ib and type-Is synaptic terminals on muscle 6/7 of A2 hemisegment for their characteristic synaptic patterning. To normalize any variation in our observations we considered several measures. Firstly, healthy and viable larvae were selected for analysis. Secondly, computer-assisted software was used to automatically measure the length of the HRP positive synaptic arbors. Thirdly, each NMJ measurements were normalized to the surface area of their corresponding muscle. Lastly, sufficient samples and controls were included to keep statistical stringency. As shown in Figure 1C-E; and quantified in Figure 1I-K, the normalized total length of *elav-GAL4* driven (CGG)_60_-EGFP motor neuron terminals showed a modest reduction (23%) with respect to controls (5.219 ± 0.252 versus 6.747 ± 0.212 × 10^3^μm; Mean ± SEM; p < 0.05, n = 16). The decrease in the synaptic span was accompanied by a 36 % escalation in the size of presynaptic varicosities (9.34 ± 0.50 versus 12.7 ± 0.41 μm2; Mean ± SEM; p < 0.05). Interestingly, however, restricted expression of (CGG)_90_–EGFP in motor neurons using *ok6-GAL4* driver lead to more severe consequences (Figure 1F-H; quantified in Figure 1I-K). In these larvae, the normalized total NMJ length decreased by 30% with respect to controls (7.25 ± 0.24 versus 5.07 ±0.18 × 10^3^μm; Mean ± SEM; p<0.0001), normalized bouton number was decreased by approximately 41% (1.29 ± 0.06 versus 0.76 ± 0.03 × 10^3^; Mean ± SEM; Bouton number/ Muscle Area (#/μm^2^)× 10^3^; p < 0.0001) and size of presynaptic varicosities escalated more than two times (8.48 ± 0.83 versus 20.1 ± 1.25; Mean ± SEM; p < 0.0001). On the contrary expression of EGFP or CGG repeats alone did not produce any of these phenotypic effects (Figure 1C-D, F; quantified in Figure 1I-K). Also, no phenotype was observed in any transgenic line when the expression was targeted to muscle cells using the *mhc-GAL4* driver, despite the fact (CGG)_90_-EGFP expression induces lethality in them at the pupal stage (quantified in Figure 1I-K). Taken together, we demonstrated that fragile X rCGG repeats exacerbate larval NMJ morphology specifically when expressed in neurons.

### Neuronal Expression of Fragile X Premutation riboCGG (rCGG) Repeats Affects the Remodeling of Synapses during Larval Development

Because boutons are continuously being added or selectively eliminated during their inception at embryonic to the late third instar larval development, we asked at what stage of development rCGG repeat specifically aggravates NMJ. The *UAS/GAL4* binary system allowed us to conditionally modulate the transgene expression level by shifting fly cultures to different temperatures (Jin et al. 2003). We performed crosses of UAS-(CGG)_90_-EGFP/TM2 Cyo with the *ok6-GAL4* driver at 18°C and continued this temperature until wandering larvae carrying genotype *(ok6-GAL4*/(CGG)_90_-EGFP) were ready for dissection. No abnormalities in average synaptic growth and bouton number were identified in them with respect to controls (1.334 ± 0.058 versus 1.284 ± 0.0496 × 10^3^μm; Mean ± SEM; Bouton number/ Muscle Area (#/μm^2^)× 10^3^). However, when the same progenies *(ok6-GAL4*/(CGG)_90_-EGFP) were shifted to 25°C post embryogenesis to increase transgene expression level, shorter synapses with engorged boutons similar to the phenotypes shown in Figure 1H were observed (1.163 ± 0.072 versus 0.755 ± 0.035 × 10^3^μm; Mean ± SEM; Bouton number/ Muscle Area (#/μm^2^)× 10^3^; p < 0.0001) (Figure 2A, quantification). On the contrary, the same crosses acclimatized to lay embryos at 25°C, and progenies shifted to 18°C post embryogenesis, did not produce any NMJ phenotype in third-instar larvae. These results suggest that the phenotype that we observed upon fragile X PM rCGG repeat commence during larval development. Importantly, under similar treatments, flies expressing EGFP only did not show such abnormalities.

**Figure 2:**
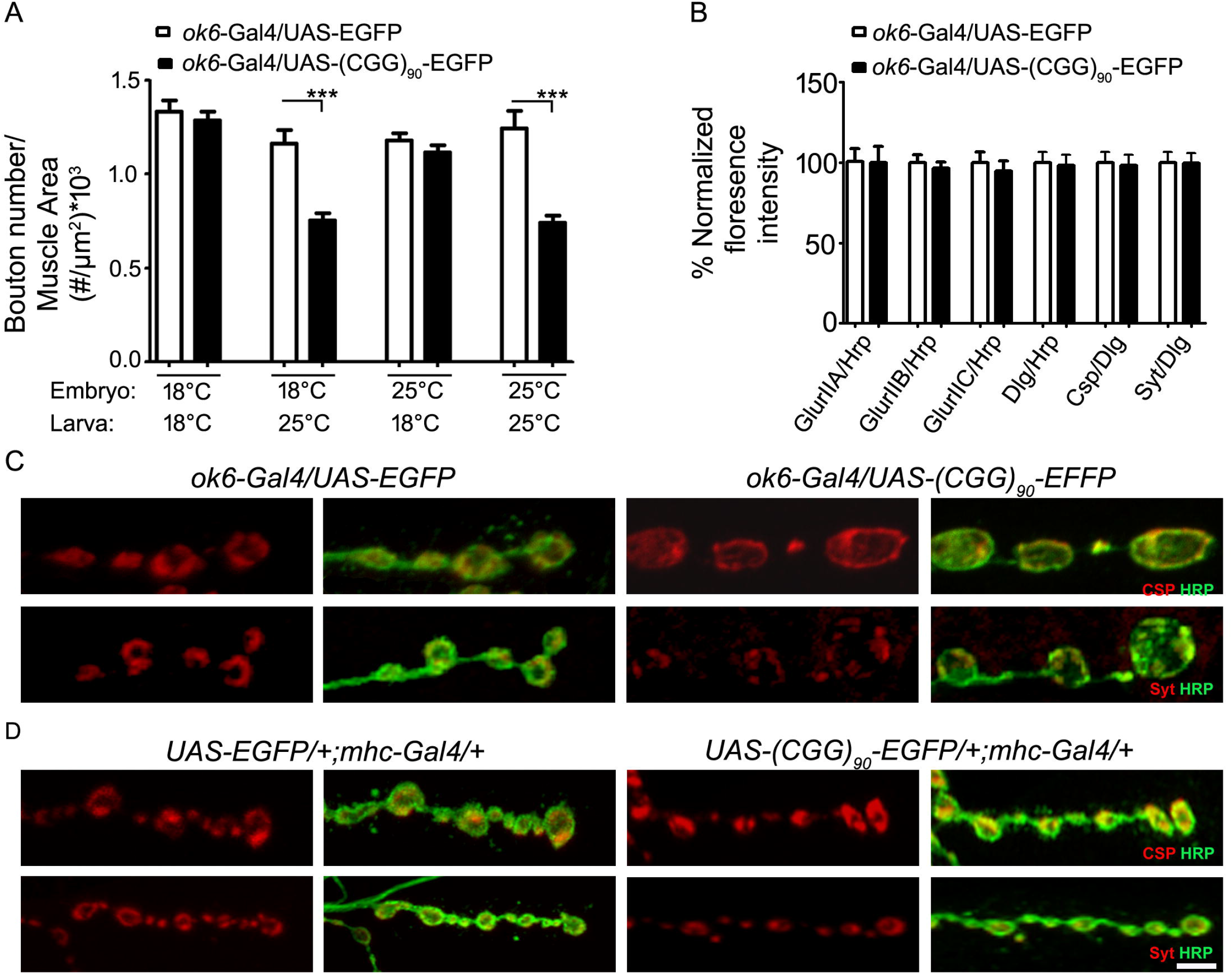
Premutation rCGG repeats in neurons affects synapse remodeling. (A) Histograms show the number of boutons per unit muscle area in *ok6-GAL4*/UAS-EGFP, and *ok6-GAL4*/UAS-(CGG)_90_-EGFP at 18°C and 25 °C. *** represents p<0.0001 (B) Histograms show average fluorescence intensity of various glutamate receptor subunits (GlurIIA; GlurIIB; GlurIIC), Subsynaptic reticulum (Dlg), CSP, Syt normalized to either HRP or Dlg taken from 4 boutons per NMJ as indicated in *ok6*-*GAL4*/UAS-EGFP; and *ok6-GAL4*/UAS-(CGG)_90_-EGFP at 25 °C. Numerical values show the number of boutons used for quantification. Student’s t-test was used for one to one analysis and One-way ANOVA with Tukey post-hoc for multiple comparisons. Representative confocal images showing type Ib boutons of muscle 4 in (C) *ok6-GAL4/*/UAS-(CGG)90-EGFP and (D) UAS-(CGG)90-EGFP/+;*mhc-Gal4*/+ at 25 °C immunolabelled with CSP or Syt shown in red, and Hrp in green. Note the uneven labeling of CSP and Syt. Scale bar represents 5μm in each case.

Many transient structures have usually been used to characterize the stages of synapse development. These include ‘ghost boutons’, presynaptic retractions (footprint), and satellite boutons. These structures are characterized and differentiated based on the presence or absence of particular molecular markers (Menon et al. 2013). We performed co-immunolabelling on NMJ fillets to reveal both pre- and post-synaptic elements of synapses simultaneously. Most of the boutons on muscle 4 or 6 of *elav-GAL4* driven (CGG)_60_-EGFP repeats were marked by simultaneous retainment of both postsynaptic (Disclarge or glutamate), and presynaptic (HRP, synaptotagmin or CSP) structures. Therefore, no signs of postsynaptic densities lacking the presynaptic component or synaptic retractions were noticed. In accord, normalization of fluorescence intensities from the same bouton showed the intactness of synapses (Figure 2B, quantification). To check the specificity of this phenotype we used a strong expression line of (CGG)_90_-EGFP using *Ok6-GAL4* driver. Interestingly, however, presynaptic expression of r(CGG)_90_-EGFP showed a high percentage of boutons having aberrant and uneven localization of vesicle markers including cysteine string protein (CSP) and synaptotagmin (Figure 2C). In normal NMJ (*ok6*-Gal4/(CGG)90-EGFP), both CSP and synaptotagmin show a ring-like localization along the periphery of the boutons (Figure 2C). No such abnormalities were detected when the expression was driven with the *mhc-GAL4* driver (Figure 2D). This observation corroborates our observations that fragile X PM rCGG repeat has severe consequences when driven presynaptically.

### Neuronal Expression of Fragile X Premutation rCGG repeats Reduces the Density of Bruchpilot (Brp) Protein at Synapses

Boutons contain multiple active zones (neurotransmitter release sites). Therefore, we asked whether altered vesicle markers or an increase in bouton size are associated with corresponding changes in active zones. Because *ok6*-*GAL4* driven (CGG)90-EGFP transgene produce prominent defects, we labeled them with active zone-specific antibody NC82 that recognizes the Bruchpilot protein (BRP). BRP is a key component of the presynaptic active zone required for active zone assembly and vesicle release (Fouquet et al. 2009, Wagh et al. 2006, Matkovic et al. 2013). As shown in Figure 3A and quantified in Figure 3D, whole synapse analysis on muscle 4 clearly shows that r(CGG)_90_ led to a remarkable reduction in the density of BRP. However, this decrease in BRP could be a consequence of strong morphological defects associated with them. Therefore, we sought to analyze such defects under moderate expression of rCGG repeats. This was achieved by using either (CGG)_60_-EGFP transgene (Figure 3B, quantified in Figure 3D) or growing transgene carrying a single copy of (CGG)_90_-EGFP repeats at 18°C (Figure 3C). Interestingly, in both cases, a significant reduction in the density of the average number of BRP punctae was noticed while maintaining morphological integrity. The reduction in BRP density seemed to be specific to NMJ as analysis of larval segmental nerves and ventral nerve cord showed uniform level and distribution of BRP punctae with no signs of aggregation (Figure 3E and 3F, quantified in Figure 3G). Also, in comparison to controls, the total amount of BRP in lysates prepared from the larval brains expressing CGG repeats remained unchanged (Figure 3H). Because active zones are normally apposed to a Glutamate Receptors (GluR) cluster (Menon et al. 2013), we asked whether the reduction in BRP punctae has some repercussions on them. Therefore, we analyzed the levels of DGluRIIC, a common subunit in ionotropic glutamate receptors. Although, most of the BRP punctae appeared normally opposed by glutamate receptors, however, some of the BRP punctae did not precede DGluRIIC labeling (Figure 3I). These results corroborate our above result that implies that active zone formation proceeds normally while subsequent synaptic remodeling events may cause a significant drop in their density. No such phenotype was observed in any transgenic line when the expression was driven with the postsynaptic *mhc-GAL4* driver.

**Figure 3:**
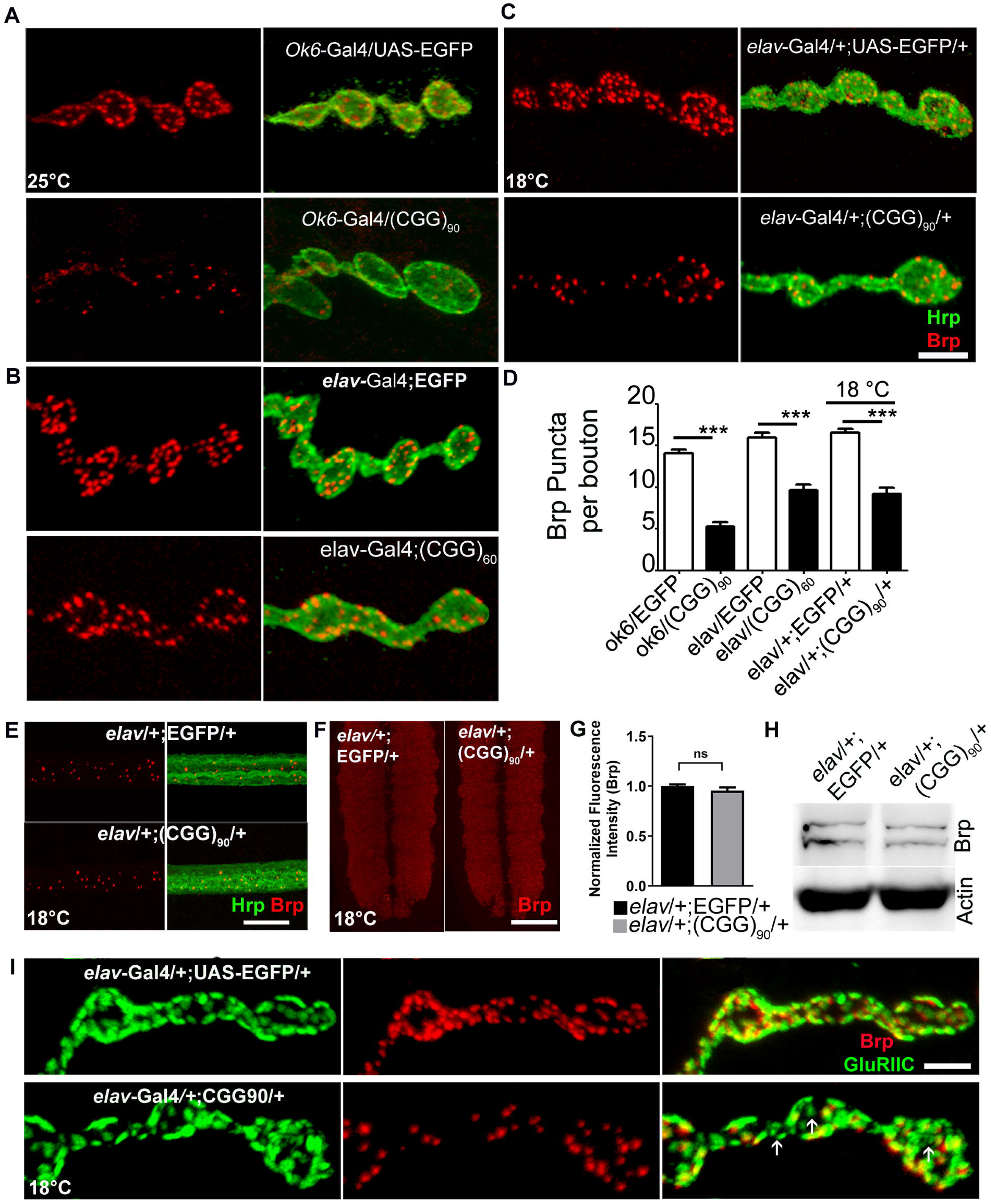
Expression of rCGG repeats decreases the number of presynaptic active zones. **(A-C)** Representative images showing NMJs of muscle 4 of (A) *ok6*-*GAL4*/UAS-EGFP and *ok6*-*GAL4*/UAS-(CGG)90-EGFP at 25 °C (B) *elav-GAL4;* UAS-EGFP and *elav-GAL4;* UAS-(CGG)60-EGFP (C) *elav*-Gal4/+; UAS-EGFP/+ and *elav-GAL4/+;* UAS-(CGG)90-EGFP/+ at 18 C. Brp is shown in red and Hrp in green. Scale bar represents 5μm. (D) Histogram plotting of Brp puncta per bouton of indicated genotypes. (E) Images showing Brp puncta in red in the segmental nerves of third instar larvae shown in green of *elav-GAL4/+;* UAS-EGFP/+ and *elav-GAL4 /*+; UAS-(CGG)_90_-EGFP/+ at 18°C. Scale bar represents 40μm. (F) Confocal images of larval third instar ventral nerve cords of *elav-GAL4/+;* UAS-EGFP/+ and *elav-GAL4 /*+; UAS-(CGG)_90_-EGFP/+ at 18°C. Red represents Brp. Scale bar represents 40μm. (G) Fluorescence intensity of Brp normalized to α-adaptin in the ventral nerve cord of the mentioned genotypes. *** represents p<0.0001. Error bars represent mean ± s.e.m. Student’s t-test was used for one to one analysis and One-way ANOVA with Tukey post-hoc for multiple comparisons. (H) Western blot showing Brp protein levels in the adult brains of *elav-GAL4/+;* UAS-EGFP/+ and *elav-GAL4 /*+; UAS-(CGG)_90_-EGFP/+ flies at 18°C. (I) Representative images showing type Ib boutons of muscle 4 overexpressing rCGG repeats immunolabelled with DGluRIII shown in green and Brp in red of *elav-GAL4/+;* UAS-EGFP/+ and *elav-GAL4 /*+; UAS-(CGG)_90_-EGFP/+ at 18°C. Note the absence of active zones opposite to many glutamate receptor fields. Scale bar represents 3μm

### Fragile X Premutation riboCGG (rCGG) Repeats induce Defects in Synaptic Transmission that Correlate with the alteration in Synapse Morphology

Structural changes in synapses are often accompanied by altered synaptic function, so we examined the consequences of CGG expression on synaptic transmission at the NMJs (Zhang and Stewart 2010). We performed intracellular electrophysiological analyses on muscle 6/7 of A2 hemisegment of third instar larval body walls. Both moderate (CGG)60 and strong (CGG)90 repeats (at 18°C) in comparison to controls exhibited a significant reduction in the nerve transmission on the arrival of impulses as was evident by a decrease in evoked excitatory junction potentials (EJP) amplitude (Figure 4A-F). EJP amplitude was reduced by 40% and 46% in r(CGG)90/+ and in homozygous r(CGG)60 expressing larvae. Reduction in EJP amplitude is often a result of reduced quantal size (post-synaptic response to single synaptic vesicle fusions) represented by miniature excitatory junction potential (mEJP) amplitude, quantal content (total synaptic vesicles fused per EJP), or a glitch in both. Assessment of mEJPs showed no reduction in r(CGG)_90_/+ expressing larvae excluding the role of quantal size in the defective synaptic transmission. However, quantal content was reduced by 53% in r(CGG)_90_/+ and 46% in r(CGG)60 expressing larvae indicating that synaptic transmission is perturbed by a decrease in the number of vesicles fusing per EJP (Figure 4J). This is consistent with our earlier observations where we found the number of vesicle releasing sites severely reduced upon expression of PM r(CGG) repeats. Interestingly, we saw a significant surge in the mEJP frequency with a nearly 40% increase in r(CGG)90/+ and 50% in r(CGG)60 expressing larvae. The mEJPs are a representative of the basal synaptic transmission indicative of homeostatic feedback mechanism, which functions when post-synapse receives less input from the pre-synapse upon PM repeats expression. As aforementioned, the glutamate receptor levels were unchanged upon rCGG expression consistent with the unrepressed quantal size despite severely impaired release machinery. This endorses that receptor clusters are normal and synaptic perturbations caused by r(CGG) repeats are not mediated through these post-synaptic components. Therefore, electrophysiological observations show that reduced release sites lead to a decrease in the release of synaptic vesicles causing altered synaptic transmission.

**Figure 4:**
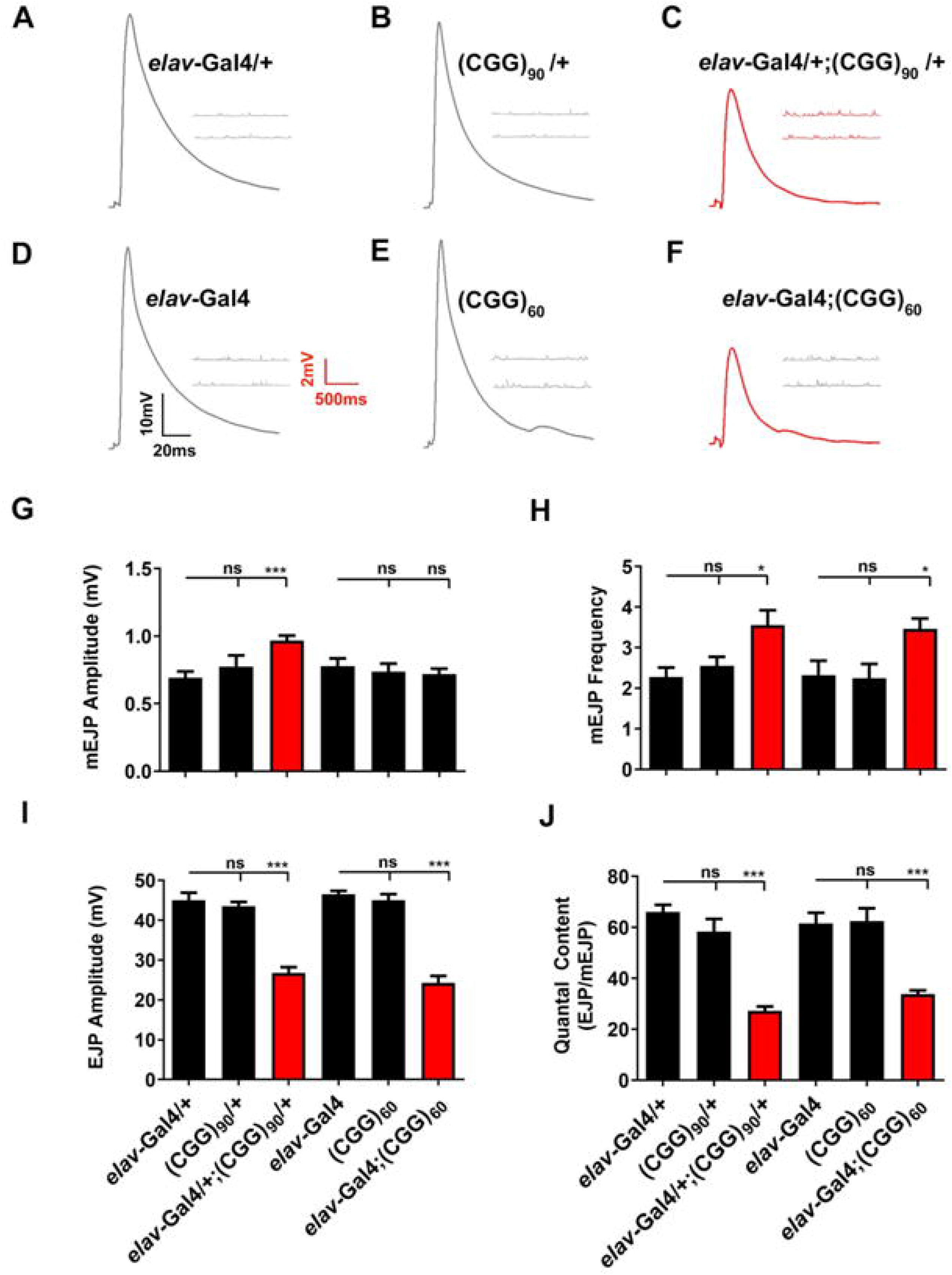
Synaptic transmission is severely affected by the expression of premutation length CGG repeats. **(A-F)** Representative evoked junction potential and miniature junction potential traces of (A) *elav*-Gal4/+; UAS-EGFP/+ (B) UAS-(CGG)90-EGFP/+ and (C) *elav*-Gal4/+; UAS-(CGG)90-EGFP/+ at 18°C. (D) *elav*-Gal4; UAS-EGFP. (E) UAS-(CGG)60-EGFP (F) *elav*-Gal4; UAS-(CGG)_60_-EGFP at 25°C. (G-J) Histograms show (G) mEJP amplitude, (H) mEJP frequency, (I) EJP amplitude, and (J) quantal content in the indicated genotypes. Error bars represent standard error of mean (SEM). * represents p<0.01, *** represents p<0.0001. One-way ANOVA followed by post hoc Tukey’s multiple comparison test was used for statistical analysis.

## Discussion

Fragile X-associated tremor/ataxia syndrome (FXTAS) is a late-onset neurodegenerative disorder and is distinct from the neurodevelopmental fragile X syndrome caused by *FMR1* gene silencing (Mila et al. 2018, Salcedo-Arellano et al. 2020). The neurological phenotypes in FXTAS patients, stemming likely from the RNA toxicity and or repeat-associated non-AUG (RAN) mediated protein toxicity, usually appear after the fifth decade of life (Swinnen, Robberecht and Van Den Bosch 2020, Todd et al. 2013, Jin et al. 2003, Jin et al. 2007). However, increasing clinical data has broadened the pathological foray of FXTAS. Several neurodevelopmental and psychological deficits have been observed in FXTAS patients, and can thus no longer recognized as a pure neurodegenerative disorder (Farzin et al. 2006, Bailey et al. 2008, Cordeiro et al. 2015, Hagerman 2012, Hagerman, Hoem and Hagerman 2010). How expanded rCGG repeat mediated pathogenesis embarks on the neurodevelopmental processes and causes misregulation is not well understood. Hence, it is imperative to trace the early pathological abnormalities caused by fragile X PM rCGG repeat. Here, using a transgenic *Drosophila* model, we spatiotemporally tracked the early events of rCGG-mediated pathogenesis in neurons. We show that the early assaults led by the expanded CGG repeats have a deleterious influence on the structure and function of the synapses in vivo. Analysis of NMJ parameters showed reduced synaptic span and larger boutons than controls (Figure 1C-H, 1I-K, quantification) that shows the expanded repeats can potentially cause cell-autonomous defects in motor neurons. The sub-synaptic analysis also revealed defects in the underlying architecture with a reduced density of presynaptic Bruchpilot protein (BRP). BRP has sequence homology to ELKS/CAST and is a key component of presynaptic active zones (neurotransmitter release sites) (Wagh et al. 2006). In flies, disruption of its action alters active zones and may leave GluR orphan without any preceding active zones (Fouquet et al. 2009). Although, in rCGG motor neurons, BRP appose normally to their GluR receptor fields, however, some of the BRP did not proceed DGluRIIC labeling (Figure 3I). These defects could be attributed to the decline in the active zones number. However, normal BRP expression levels and delivery of BRP to the synapses implies that active zone formation proceeds normally while subsequent synaptic remodeling events may be compromised. Significantly, this manifests in the strong reduction in the quantal content, a measure of total synaptic vesicles released per excitation potential (Figure 4J). This endorses that receptor clusters are normal, and synapse perturbation caused by r(CGG) repeats is not mediated through these postsynaptic compartments. This is evidently due to the reduction in active zones known to modulate synaptic communication (Ehmann et al. 2018).

Synaptic defects are common in early appearing neurological diseases, including Fragile X syndrome, autism spectrum disorders (ASD), schizophrenia (SCZ), and bipolar disorder (BP)(Hagerman 2012, Chelly et al. 2006, Parikshak et al. 2013, Parikshak, Gandal and Geschwind 2015, Bagni and Greenough 2005). On the other hand, in late-onset neurodegeneration, such as Alzheimer’s (AD), Parkinson’s (PD), and Huntington’s (HD) diseases, cell death is the inevitable end-result of an ongoing pathophysiological cascade. These diseases also converge on to the same structural and pathological features like mitochondrial abnormalities (Haelterman et al. 2014, Wang, Berry-Kravis and Hagerman 2010), synaptic defects (Bendor, Logan and Edwards 2013, Spires-Jones and Hyman 2014), and in some cases cause the absence of assortment of presynaptic proteins like active zone components and vesicle-associated proteins(Scott et al. 2010, Chouhan et al. 2016, Jack et al. 2013, Bendor et al. 2013, Spires-Jones and Hyman 2014). PM repeats cause early defects in neuronal morphology (Chen et al. 2010, Qin et al. 2011), spontaneous calcium oscillations, and clustered burst firing in the animal models (Cao et al. 2012). Solidifying these observations is the growing clinical data showing alterations in the brain well before the onset of FXTAS. These include a drop in cerebellar and brain stem size volume (Hashimoto et al. 2011), white matter tract changes (Wang et al. 2013), and functional MRI recordings (Wang et al. 2012). Thus, it is evident that comorbid neurodevelopmental conditions with FXTAS are growing; however, the relationship between their manifestations remains obscure. Our findings presented here suggest rCGG-repeats cause synaptic abnormalities. Whether these abnormalities proceed neuronal cell death, as seen in FXTAS, remains to be investigated. However, these abnormalities could provide a better understanding of the underlying shared mechanisms among comorbid neurological disorders. Such information will provide lead to improved disease biomarkers and novel therapeutic approaches. In summary, our observation in a transgenic model of fragile X PM rCGG repeats shows that the structural and functional alterations in synapses caused by fragile X PM rCGG repeats may contribute to FXTAS pathogenesis, and mechanisms could be via BRP mediated impact on active zone seen upon fragile X PM rCGG expression.

## Experimental Procedures

### Fly stocks and genetics

All the fly strains were reared on a standard cornmeal media at 25°C unless otherwise mentioned. Transgenic flies with both moderate (pUAST-(CGG)_60_-EGFP), or strong (pUAST-(CGG)_90_-EGFP) transgene expressions and control (pUAST-EGFP) used in this study were generated previously (Jin et al., 2003). The presence of CGG repeats in the transgenic flies was confirmed by polymerase chain reaction (PCR) using C and F primers as described previously (Fu et al., 1991,Tassone et al., 2008). The expression of these repeats was checked by reverse transcription PCR using primers against the eGFP tag (Todd et al., 2010). *GAL-4* stocks *elav-GAL4,* and *ok6-GAL4* were procured from the Bloomington Drosophila Stock Center, Indiana University.

### Immunohistochemistry and antibodies

For NMJ immunohistochemistry wandering third instar larvae were dissected in ice-cold calcium-free hemolymph-like saline (HL3) solution and fixed in 4% paraformaldehyde in 1X phosphate-buffered saline (PBS), pH 7.2 for 30 minutes or in Bouin’s fixative (Glutamate receptor staining) for 5 minutes. After washing with PBS, containing 0.2% Triton X-100 (PBT), larval fillets were blocked in 5% bovine serum albumin (BSA) in PBT for 1hr and then incubated overnight with the primary antibody in PBT at 4°C. The larvae were then washed in PBT and incubated in secondary antibody in PBT for 1hr followed by washing steps and mounted in VectaShield (Vector Laboratories, Burlingame, CA) on slides. The following monoclonal antibodies from Developmental Studies Hybridoma Bank (University of Iowa, USA) were used for immunostaining: anti-CSP (1:50), anti-Brp^Nc82^(1:50), and anti-GluRIIA (1:50). DGluRIIB (1:1000) and DGluRIII (1:5000) were provided by Aaron DiAntonio(Marrus et al., 2004). Anti-α-adaptin (Choudhury et al. 2016) was used as an internal control for Brp staining in larval brains. Secondary antibodies conjugated to fluorophores Alexa Fluor 488, Alexa Fluor 568, Alexa Fluor 555, and Alexa Fluor 633 (Molecular Probes, Thermo Fischer Scientific) were used at 1:800 dilution. Anti-HRP antibodies conjugated to Alexa-488 or Rhodamine were used at 1:1000 dilution.

### Larval crawling assay

The third instar larvae were gently collected and washed clean with deionized water. The larvae were subsequently transferred on a 2% agarose gel in a transparent Petri dish placed on top of a gridline marked surface. The larvae were allowed to acclimatize for some time before making quantifications. The average distance traversed by larvae in 60 sec was calculated for each genotype. The experiments were performed in uncrowded conditions. Statistical analyses based on Students t-test were performed using GraphPad Prism software (GraphPad Software, San Diego)

### Electrophysiology

Intracellular electrophysiological recordings were performed at room temperature. Glass microelectrodes filled with 3M KCl with resistance between 12-20Ω were used. Experiments were performed on third instar larval NMJs (muscle 6, A2 hemisegment) in HL3 saline with composition as (in mM): NaCl 70, KCl 5, MgCl_2_ 20, NaHCO3 10, sucrose 115, trehalose 5, and HEPES 5 with pH 7.2 supplemented with 1.5mM calcium. Recordings were done from muscles having resting potential between −60mV to −75mV and input resistance of ≥4MΩ. For recording EJPs, nerves were stimulated at 1Hz and spontaneous release events lasted for the 60s. For synaptic depression experiments, nerves were subjected to 10Hz stimulation for 5 minutes. To test the recovery, nerves were stimulated at 10Hz for 5 minutes followed by 90s rest, and then test stimulated every 30s. Estimation of quantal content was achieved by the ratio of average EJP over average mEJP amplitude for each NMJ. The signals were amplified using Axoclamp 900A and data acquisition using Digidata 1440A and pClamp10 software (Axon Instruments, Molecular Devices, USA). Mini-Analysis (Synaptosoft) was used for data analysis.

### Western blotting

Third instar larval brains were manually homogenized in 1.5ml Eppendorf tubes at 65°C using a micropestle in an equal amount of 2X Laemmli buffer and the samples were heated at 95°C for 10 min. The debris was pelleted by centrifugation at 13000×g for 5 min and the protein sample equivalent to 5 heads was resolved on 8% denaturing SDS-PAGE. Proteins were transferred to the Hybond-PVDF-LFP membrane (Amersham, GE Healthcare Life Sciences) and blocked with 5% skimmed milk in 1X Tris-buffered saline, supplemented with 0.1% Tween-20 (TBST) for 1hr at room temperature. The membrane was probed with primary antibodies against Brp^Nc82^ (1:500) and α-Actin (Cell signaling) in 1X TBST containing 2% BSA at 4°C overnight followed by washing steps in TBST. This was followed by incubation with secondary antibodies conjugated to HRP in 1X TBST, (1:10000) for 1hr followed by washing steps. The membranes were incubated with ECL Prime Western Blotting Detection Reagent (GE Healthcare) and signals were acquired using Fuji LAS-4000 Image Analyzer (Fuji Film, Tokyo, Japan). The images were analyzed using Image Gauge software (Fuji Film).

### Image acquisition and analysis

Images were captured using a Zeiss LSM780 confocal microscope. NMJ morphometric analyses were performed at muscle 6/7 of abdominal segment 2 except for Brp puncta quantification which was done at muscle 4 of abdominal segment 2 for convenience. Neuronal membranes were labeled with anti-HRP antibody and boutons were specifically labeled with vesicle specific anti-CSP antibody. Images for fluorescence quantifications were acquired with the same parameters and settings. Fluorescence intensities were quantified using Image J (National Institute of Health) or Zen software (Carl Zeiss). The data are reported as mean ±SEM. Images were processed by Adobe Photoshop 7.0 (Adobe Systems, San Jose, CA). For statistical analysis a two-tailed Student’s t-test was used for experiments involving two data sets otherwise one-way ANOVA with Tukey’s multiple comparison post-test was employed. The asterisks in the figures represent the level of significance: *, P≤0.05; **, P≤0.01 and ***, P≤ 0.001.

## Authors’ contributions

SAB carried out experiments. SAB and ZM carried out the electrophysiology experiments. AY helped out in experiments. AQ and SAB carried out the phenotypic analysis. AQ, AY and VK participated in coordinating the study. SAB and Abrar Q prepared the figures. Abrar Q conceived the study and wrote the final version of the manuscript. All authors have read and approved the final version of the manuscript.

## Acknowledgements

We are especially grateful to Peng Jin for providing tools required for the study (transgenic flies carrying PM CGG repeats, and controls). We are thankful to Aaron DiAntonio, the Developmental Studies Hybridoma Bank, and Bloomington Stock Centers for providing antibodies and important fly lines. We thank confocal facility at IISER Bhopal for help with microscopy used in this study. This work was supported by the Department of Biotechnology, Govt of India (BT/RLF/Re-entry/51/2012), and UGC-BSR grant to Abrar Q. SAB was supported by CSIR, New Delhi through research fellowship.

## References

Alvarez-Mora, M. I., L. Rodriguez-Revenga, I. Madrigal, M. Guitart-Mampel, G. Garrabou & M. Mila (2017) Impaired Mitochondrial Function and Dynamics in the Pathogenesis of FXTAS. Mol Neurobiol, 54, 6896–6902.

Ariza, J., H. Rogers, A. Monterrubio, A. Reyes-Miranda, P. J. Hagerman & V. Martinez-Cerdeno (2016) A Majority of FXTAS Cases Present with Intranuclear Inclusions Within Purkinje Cells. Cerebellum, 15, 546–51.

Bagni, C. & W. T. Greenough (2005) From mRNP trafficking to spine dysmorphogenesis: the roots of fragile X syndrome. Nat Rev Neurosci, 6, 376–87.

Bailey, D. B., M. Raspa, M. Olmsted & D. B. Holiday (2008) Co-occurring conditions associated with FMR1 gene variations: findings from a national parent survey. Am J Med Genet A, 146A, 2060–9.

Bendor, J. T., T. P. Logan & R. H. Edwards (2013) The function of α-synuclein. Neuron, 79, 1044–66.

Berman, R. F. & R. Willemsen (2009) Mouse models of fragile X-associated tremor ataxia. J Investig Med, 57, 837–41.

Cao, Z., S. Hulsizer, F. Tassone, H. T. Tang, R. J. Hagerman, M. A. Rogawski, P. J. Hagerman & I. N. Pessah (2012) Clustered burst firing in FMR1 premutation hippocampal neurons: amelioration with allopregnanolone. Hum Mol Genet, 21, 2923–35.

Chelly, J., M. Khelfaoui, F. Francis, B. Chérif & T. Bienvenu (2006) Genetics and pathophysiology of mental retardation. Eur J Hum Genet, 14, 701–13.

Chen, Y., F. Tassone, R. F. Berman, P. J. Hagerman, R. J. Hagerman, R. Willemsen & I. N. Pessah (2010) Murine hippocampal neurons expressing Fmr1 gene premutations show early developmental deficits and late degeneration. Hum Mol Genet, 19, 196–208.

Chonchaiya, W., J. Au, A. Schneider, D. Hessl, S. W. Harris, M. Laird, Y. Mu, F. Tassone, D. V. Nguyen & R. J. Hagerman (2012) Increased prevalence of seizures in boys who were probands with the FMR1 premutation and co-morbid autism spectrum disorder. Hum Genet, 131, 581–9.

Choudhury, S. D., Z. Mushtaq, S. Reddy-Alla, S. S. Balakrishnan, R. S. Thakur, K. S. Krishnan, P. Raghu, M. Ramaswami & V. Kumar (2016) σ2-Adaptin Facilitates Basal Synaptic Transmission and Is Required for Regenerating Endo-Exo Cycling Pool Under High-Frequency Nerve Stimulation in Drosophila. Genetics, 203, 369–85.

Chouhan, A. K., C. Guo, Y. C. Hsieh, H. Ye, M. Senturk, Z. Zuo, Y. Li, S. Chatterjee, J. Botas, G. R. Jackson, H. J. Bellen & J. M. Shulman (2016) Uncoupling neuronal death and dysfunction in Drosophila models of neurodegenerative disease. Acta Neuropathol Commun, 4, 62.

Colak, D., N. Zaninovic, M. S. Cohen, Z. Rosenwaks, W. Y. Yang, J. Gerhardt, M. D. Disney & S. R. Jaffrey (2014) Promoter-bound trinucleotide repeat mRNA drives epigenetic silencing in fragile X syndrome. Science, 343, 1002–5.

Cordeiro, L, F. Abucayan, R. Hagerman, F. Tassone & D. Hessl (2015) Anxiety disorders in fragile X premutation carriers: Preliminary characterization of probands and non-probands. Intractable Rare Dis Res, 4, 123–30.

Dombrowski, C., S. Lévesque, M. L. Morel, P. Rouillard, K. Morgan & F. Rousseau (2002) Premutation and intermediate-size FMR1 alleles in 10572 males from the general population: loss of an AGG interruption is a late event in the generation of fragile X syndrome alleles. Hum Mol Genet, 11, 371–8.

Ehmann, N., D. Owald & R. J. Kittel (2018) Drosophila active zones: From molecules to behaviour. Neurosci Res, 127, 14–24.

Farzin, F., H. Perry, D. Hessl, D. Loesch, J. Cohen, S. Bacalman, L. Gane, F. Tassone, P. Hagerman & R. Hagerman (2006) Autism spectrum disorders and attention-deficit/hyperactivity disorder in boys with the fragile X premutation. J Dev Behav Pediatr, 27, S137–44.

Fouquet, W., D. Owald, C. Wichmann, S. Mertel, H. Depner, M. Dyba, S. Hallermann, R. J. Kittel, S. Eimer & S. J. Sigrist (2009) Maturation of active zone assembly by Drosophila Bruchpilot. J Cell Biol, 186, 129–45.

Garcia-Arocena, D., J. E. Yang, J. R. Brouwer, F. Tassone, C. Iwahashi, E. M. Berry-Kravis, C. G. Goetz, A. M. Sumis, L. Zhou, D. V. Nguyen, L. Campos, E. Howell, A. Ludwig, C. Greco, R. Willemsen, R. J. Hagerman & P. J. Hagerman (2010) Fibroblast phenotype in male carriers of FMR1 premutation alleles. Hum Mol Genet, 19, 299–312.

Goodlin-Jones, B. L., F. Tassone, L. W. Gane & R. J. Hagerman (2004) Autistic spectrum disorder and the fragile X premutation. J Dev Behav Pediatr, 25, 392–8.

Goodrich-Hunsaker, N. J., L. M. Wong, Y. McLennan, S. Srivastava, F. Tassone, D. Harvey, S. M. Rivera & T. J. Simon (2011) Young adult female fragile X premutation carriers show age- and genetically-modulated cognitive impairments. Brain Cogn, 75, 255–60.

Haelterman, N. A., W. H. Yoon, H. Sandoval, M. Jaiswal, J. M. Shulman & H. J. Bellen (2014) A mitocentric view of Parkinson’s disease. Annu Rev Neurosci, 37, 137–59.

Hagerman, P. (2013) Fragile X-associated tremor/ataxia syndrome (FXTAS): pathology and mechanisms. Acta Neuropathol, 126, 1–19.

Hagerman, P. J. (2012) Current Gaps in Understanding the Molecular Basis of FXTAS. Tremor Other Hyperkinet Mov(NY), 2.

Hagerman, P. J. & R. J. Hagerman (2015) Fragile X-associated tremor/ataxia syndrome. Ann N Y Acad Sci, 1338, 58–70.

Hagerman, R., G. Hoem & P. Hagerman (2010) Fragile X and autism: Intertwined at the molecular level leading to targeted treatments. Mol Autism, 1, 12.

Hagerman, R. J. & P. J. Hagerman (2002) The fragile X premutation: into the phenotypic fold. Curr Opin Genet Dev, 12, 278–83.

Hagerman, R. J., L. W. Staley, R. O’Conner, K. Lugenbeel, D. Nelson, S. D. McLean & A. Taylor (1996) Learning-disabled males with a fragile X CGG expansion in the upper premutation size range. Pediatrics, 97, 122–6.

Hashimoto, R., A. K. Javan, F. Tassone, R. J. Hagerman & S. M. Rivera (2011) A voxel-based morphometry study of grey matter loss in fragile X-associated tremor/ataxia syndrome. Brain, 134, 863–78.

Hunsaker, M. R., H. J. Wenzel, R. Willemsen & R. F. Berman (2009) Progressive spatial processing deficits in a mouse model of the fragile X premutation. Behav Neurosci, 123, 1315–24.

Jack, C. R., D. S. Knopman, W. J. Jagust, R. C. Petersen, M. W. Weiner, P. S. Aisen, L. M. Shaw, P. Vemuri, H. J. Wiste, S. D. Weigand, T. G. Lesnick, V. S. Pankratz, M. C. Donohue & J. Q. Trojanowski (2013) Tracking pathophysiological processes in Alzheimer’s disease: an updated hypothetical model of dynamic biomarkers. Lancet Neurol, 12, 207–16.

Jacquemont, S., R. J. Hagerman, M. Leehey, J. Grigsby, L. Zhang, J. A. Brunberg, C. Greco, V. Des Portes, T. Jardini, R. Levine, E. Berry-Kravis, W. T. Brown, S. Schaeffer, J. Kissel, F. Tassone & P. J. Hagerman (2003) Fragile X premutation tremor/ataxia syndrome: molecular, clinical, and neuroimaging correlates. Am J Hum Genet, 72, 869–78.

Jacquemont, S., R. J. Hagerman, M. A. Leehey, D. A. Hall, R. A. Levine, J. A. Brunberg, L. Zhang, T. Jardini, L. W. Gane, S. W. Harris, K. Herman, J. Grigsby, C. M. Greco, E. Berry-Kravis, F. Tassone & P. J. Hagerman (2004) Penetrance of the fragile X-associated tremor/ataxia syndrome in a premutation carrier population. JAMA, 291, 460–9.

Jin, P., R. Duan, A. Qurashi, Y. Qin, D. Tian, T. C. Rosser, H. Liu, Y. Feng & S. T. Warren (2007) Pur alpha binds to rCGG repeats and modulates repeat-mediated neurodegeneration in a Drosophila model of fragile X tremor/ataxia syndrome. Neuron, 55, 556–64.

Jin, P., D. C. Zarnescu, F. Zhang, C. E. Pearson, J. C. Lucchesi, K. Moses & S. T. Warren (2003) RNA-mediated neurodegeneration caused by the fragile X premutation rCGG repeats in Drosophila. Neuron, 39, 739–47.

Kremer, E. J., M. Pritchard, M. Lynch, S. Yu, K. Holman, E. Baker, S. T. Warren, D. Schlessinger, G. R. Sutherland & R. I. Richards (1991) Mapping of DNA instability at the fragile X to a trinucleotide repeat sequence p(CCG)n. Science, 252, 1711–4.

Matkovic, T., M. Siebert, E. Knoche, H. Depner, S. Mertel, D. Owald, M. Schmidt, U. Thomas, A. Sickmann, D. Kamin, S. W. Hell, J. Bürger, C. Hollmann, T. Mielke, C. Wichmann & S. J. Sigrist (2013) The Bruchpilot cytomatrix determines the size of the readily releasable pool of synaptic vesicles. J Cell Biol, 202, 667–83.

Menon, K. P., R. A. Carrillo & K. Zinn (2013) Development and plasticity of the Drosophila larval neuromuscular junction. Wiley Interdiscip Rev Dev Biol, 2, 647–70.

Mila, M., M. I. Alvarez-Mora, I. Madrigal & L. Rodriguez-Revenga (2018) Fragile X syndrome: An overview and update of the FMR1 gene. Clin Genet, 93, 197–205.

Parikshak, N. N., M. J. Gandal & D. H. Geschwind (2015) Systems biology and gene networks in neurodevelopmental and neurodegenerative disorders. Nat Rev Genet, 16, 441–58.

Parikshak, N. N., R. Luo, A. Zhang, H. Won, J. K. Lowe, V. Chandran, S. Horvath & D. H. Geschwind (2013) Integrative functional genomic analyses implicate specific molecular pathways and circuits in autism. Cell, 155, 1008–21.

Prokop, A. (2006) Organization of the efferent system and structure of neuromuscular junctions in Drosophila. Int Rev Neurobiol, 75, 71–90.

Qin, M., A. Entezam, K. Usdin, T. Huang, Z. H. Liu, G. E. Hoffman & C. B. Smith (2011) A mouse model of the fragile X premutation: effects on behavior, dendrite morphology, and regional rates of cerebral protein synthesis. Neurobiol Dis, 42, 85–98.

Qurashi, A., W. Li, J. Y. Zhou, J. Peng & P. Jin (2011) Nuclear accumulation of stress response mRNAs contributes to the neurodegeneration caused by Fragile X premutation rCGG repeats. PLoS Genet, 7, e1002102.

Ross-Inta, C., A. Omanska-Klusek, S. Wong, C. Barrow, D. Garcia-Arocena, C. Iwahashi, E. Berry-Kravis, R. J. Hagerman, P. J. Hagerman & C. Giulivi (2010) Evidence of mitochondrial dysfunction in fragile X-associated tremor/ataxia syndrome. Biochem J, 429, 545–52.

Rousseau, F., P. Rouillard, M. L. Morel, E. W. Khandjian & K. Morgan (1995) Prevalence of carriers of premutation-size alleles of the FMR1 gene--and implications for the population genetics of the fragile X syndrome. Am J Hum Genet, 57, 1006–18.

Salcedo-Arellano, M. J., B. Dufour, Y. McLennan, V. Martinez-Cerdeno & R. Hagerman (2020) Fragile X syndrome and associated disorders: Clinical aspects and pathology. Neurobiol Dis, 136, 104740.

Scott, D. A., I. Tabarean, Y. Tang, A. Cartier, E. Masliah & S. Roy (2010) A pathologic cascade leading to synaptic dysfunction in alpha-synuclein-induced neurodegeneration. J Neurosci, 30, 8083–95.

Sellier, C., R. A. M. Buijsen, F. He, S. Natla, L. Jung, P. Tropel, A. Gaucherot, H. Jacobs, H. Meziane, A. Vincent, M. F. Champy, T. Sorg, G. Pavlovic, M. Wattenhofer-Donze, M. C. Birling, M. Oulad-Abdelghani, P. Eberling, F. Ruffenach, M. Joint, M. Anheim, V. Martinez-Cerdeno, F. Tassone, R. Willemsen, R. K. Hukema, S. Viville, C. Martinat, P. K. Todd & N. Charlet-Berguerand (2017) Translation of Expanded CGG Repeats into FMRpolyG Is Pathogenic and May Contribute to Fragile X Tremor Ataxia Syndrome. Neuron, 93, 331–347.

Spires-Jones, T. L. & B. T. Hyman (2014) The intersection of amyloid beta and tau at synapses in Alzheimer’s disease. Neuron, 82, 756–71.

Swinnen, B., W. Robberecht & L. Van Den Bosch (2020) RNA toxicity in non-coding repeat expansion disorders. Embo j, 39, e101112.

Tassone, F., R. J. Hagerman, A. K. Taylor, L. W. Gane, T. E. Godfrey & P. J. Hagerman (2000) Elevated levels of FMR1 mRNA in carrier males: a new mechanism of involvement in the fragile-X syndrome. Am J Hum Genet, 66, 6–15.

Todd, P. K., S. Y. Oh, A. Krans, F. He, C. Sellier, M. Frazer, A. J. Renoux, K. C. Chen, K. M. Scaglione, V. Basrur, K. Elenitoba-Johnson, J. P. Vonsattel, E. D. Louis, M. A. Sutton, J. P. Taylor, R. E. Mills, N. Charlet-Berguerand & H. L. Paulson (2013) CGG repeat-associated translation mediates neurodegeneration in fragile X tremor ataxia syndrome. Neuron, 78, 440–55.

Verkerk, A. J., M. Pieretti, J. S. Sutcliffe, Y. H. Fu, D. P. Kuhl, A. Pizzuti, O. Reiner, S. Richards, M. F. Victoria & F. P. Zhang (1991) Identification of a gene (FMR-1) containing a CGG repeat coincident with a breakpoint cluster region exhibiting length variation in fragile X syndrome. Cell, 65, 905–14.

Wagh, D. A., T. M. Rasse, E. Asan, A. Hofbauer, I. Schwenkert, H. Dürrbeck, S. Buchner, M. C. Dabauvalle, M. Schmidt, G. Qin, C. Wichmann, R. Kittel, S. J. Sigrist & E. Buchner (2006) Bruchpilot, a protein with homology to ELKS/CAST, is required for structural integrity and function of synaptic active zones in Drosophila. Neuron, 49, 833–44.

Wang, J. M., K. Koldewyn, R. Hashimoto, A. Schneider, L. Le, F. Tassone, K. Cheung, P. Hagerman, D. Hessl & S. M. Rivera (2012) Male carriers of the FMR1 premutation show altered hippocampal-prefrontal function during memory encoding. Front Hum Neurosci, 6, 297.

Wang, J. Y., D. Hessl, A. Schneider, F. Tassone, R. J. Hagerman & S. M. Rivera (2013) Fragile X-associated tremor/ataxia syndrome: influence of the FMR1 gene on motor fiber tracts in males with normal and premutation alleles. JAMA Neurol, 70, 1022–9.

Wang, L. W., E. Berry-Kravis & R. J. Hagerman (2010) Fragile X: leading the way for targeted treatments in autism. Neurotherapeutics, 7, 264–74.

Zhang, B. & B. Stewart (2010) Electrophysiological recording from a ‘model’ cell. Cold Spring Harb Protoc, 2010, pdb.prot5486.

